# Genetic diversity of *Legionella* species in culture-negative clinical and environmental specimens by sequencing the 23S-5S ribosomal intergenic spacer region

**DOI:** 10.64898/2026.09.18.752540

**Authors:** Camille Jacqueline, Aurélie Peticca, Justine Lannes, Matthieu Curtil-Dit-Galin, Marine Ibranosyan, Laetitia Béraud, Ghislaine Descours, Sophie Jarraud, Christophe Ginevra

## Abstract

The diagnosis of Legionnaires’ disease (LD) caused by *Legionella* non-*pneumophila* species is likely to increase with broader use of PCR targeting *Legionella* spp. In this context, accurate species identification in PCR-positive but culture-negative samples is essential to improve understanding of disease epidemiology and to support source attribution. Here, we presented a validated and user-friendly bioinformatic pipeline compatible with next-generation sequencing (NGS) for analyzing the hypervariable 23S-5S region, paired with a curated database encompassing all described *Legionella* species as of January 2026. Parameters were optimized for sensitivity and specificity using both strains and culture-positive clinical and environmental samples. We then applied the pipeline retrospectively to 92 culture-negative PCR-positive samples collected from 2023 to 2025. *Legionella* species were successfully assigned in 60% (55/92) of tested samples and revealed a high diversity. Co-infections were detected in clinical samples, including combinations of *L. pneumophila* with *L. longbeachae* or *L. bozemanii*, while environmental samples contained up to six different species. These results demonstrate that 23S-5S amplicon NGS enables species-level identification in the absence of cultured isolates, improving surveillance of non-pneumophila *Legionella* cases. The proposed pipeline, implemented in QIIME2 and accompanied by a publicly available database, provides a practical framework for routine molecular monitoring and outbreak investigation.

**Author Notes:** All supporting data have been provided within the article or through supplementary data files. Four supplementary tables are available with the online version of this article.

**Impact statement:** Legionnaires’ disease caused by non-*pneumophila Legionella* species remains under-characterized because species identification is often impossible when cultures are negative. We developed and validated a next-generation sequencing approach targeting the *Legionella* 23S-5S region, coupled with a curated reference database and a user-friendly bioinformatic pipeline, enabling species-level identification directly from PCR-positive samples. Application to clinical and environmental specimens revealed substantial *Legionella* diversity in environmental samples and detected mixed-species infections that would otherwise have been missed. This open and reproducible framework strengthens molecular surveillance and outbreak investigations in the era of culture-independent diagnostics, as well as our understanding of *Legionella* diversity.

**Data summary:** All isolates whole-genome sequences used in this work are available in Table S1, available in the online version of this article. Amplicon-specific information can be found in Table S2, available in the online version of this article.

## Introduction

Legionnaires’ disease (LD) is a severe and potentially fatal form of bacterial pneumonia caused by *Legionella* species (*Legionella* spp.), Gram-negative bacteria commonly found in fresh-water and soil environments (Rowbotham, 1980). *Legionella pneumophila* is the predominant species and accounts for the vast majority of human cases. Among the more than 60 described *Legionella* spp. (https://lpsn.dsmz.de/genus/legionella), approximatively half have been reported to infect humans (Barskey et al., 2022). Transmission typically occurs through inhalation of contaminated aerosols originating from environmental sources (Muder et al., 1986). Frequently implicated sources include water systems of large buildings, cooling towers, spa pools, and decorative fountains. In addition, several *Legionella* spp. have been detected in soils, potting mixes, and composts. In 2022, 4% of LD cases in Europe were associated with non-*pneumophila Legionella* spp. among culture-confirmed cases (ECDC, 2026). *L. longbeachae* was the most frequently isolated species after *L. pneumophila* (ECDC, 2026).

Accurate identification of the *Legionella* spp. is essential for clinical management, outbreak investigation and for implementation of efficient control measure through source attribution. However, LD caused by non-*L. pneumophila* is likely underreported for several reasons. First, most urinary antigen tests primarily detect *L. pneumophila* serogroup 1, with limited exceptions such as *L. longbeachae* detection using the IMMUVIEW® assay or *Legionella* spp. by LIAISON® assay (Diasorin, Italy). Second, culture remains technically demanding and is performed in only approximatively 10% of cases in Europe (ECDC, 2026). Third, the presence of multiple co-occurring *Legionella* spp. in environmental and clinical specimens complicates detection and isolation (Buchbinder et al., 2004; Coscollá et al., 2014; Wewalka et al., 2014). In environmental matrices such as wastewater, soil, or compost, overgrowth by competing microorganisms further reduces isolation efficiency.

Several molecular methods have been proposed to characterize *Legionella* spp. in the absence of strains, but their routine application remains limited. The 23S-5S ribosomal intergenic spacer region (ISR) has demonstrated strong discriminatory potential. Grattard et al. (2006) developed primers targeting this region by real-time PCR. Through Sanger sequencing of amplified products from respiratory samples, they were able to assign species in the absence of strains (Grattard et al., 2006). However, their study included only 10 samples with confirmed *L. pneumophila* infection, limiting broader evaluation. Subsequently, another study described a real-time PCR assay followed by amplicon Sanger sequencing capable of differentiating 50 non-*pneumophila* species (Yang et al., 2010). Although validated on a bigger yet limited dataset of 27 PCR-positive, culture-negative samples, species-level identification was achieved in 70% of cases.

Given these limitations and the development of next-generation sequencing, further exploration of 23S-5S amplicon sequencing for comprehensive *Legionella* spp. identification is warranted. We hypothesized that optimized 23S-5S amplicon NGS sequencing combined with a comprehensive curated database would enable reliable species-level identification directly from culture-negative samples in routine reference laboratories, including in the presence of multiple *Legionella* spp. In this retrospective study, we applied an optimized 23S– 5S typing approach to 92 PCR-positive, culture-negative clinical and environmental samples collected and tested between 2023 and 2025. We developed a user-friendly bioinformatic pipeline for analysis and visualization, designed for sustainable routine implementation. Additionally, we provide an updated curated reference database including all *Legionella* spp. described as of January 2026, publicly available via GitHub. The sensitivity and specificity of the pipeline was validated using a panel of 155 genomes, including 90 reference genomes, and using 20 culture-positive samples representing multiple *Legionella* spp.

## Materiel and methods

### Bacterial strains

A total of 65 *Legionella* strains provided by the French Legionella Reference Center (CNR, Lyon, France) were included for specificity assessment (Table 1). Strains were grown on buffered charcoal yeast extract (BCYE) agar (Oxoid, Dardilly, France) at 35°C under in a humidified atmosphere containing 2.5% CO_2_ for 72h prior DNA extraction.

**Table 1:** CNR-L strains included in the specificity analysis.

| Species | Number of CNR-L strains |
| --- | --- |
| *L. anisa* | 28 |
| *L. bozemanii* | 8 |
| *L. cincinnatiensis* | 2 |
| *L. dumoffii* | 2 |
| *L. gormanii* | 1 |
| *L. longbeachae* | 17 |
| *L. sainthelensi* | 2 |
| *L. steelei* | 1 |
| *L. taurinensis* | 4 |

Genomic DNA was extracted using the Maxwell® automated extraction system (Promega, Charbonnières-Les-Bains, France) according to the manufacturer’s instructions for kit RSC Blood DNA kit (reference AS1400). DNA concentration and quality were assessed using Quantus fluorometer.

To further test the specificity of the database, whole genome sequences were obtained from ENA for 90 reference strains (Supplementary Table S1).

### Clinical and environmental samples

The study included 92 PCR-positive (for *Legionella* spp. according to (Templeton et al., 2003)), culture-negative specimens collected and tested between 2023 to 2025. Among these, 59 were clinical samples (biopsy, respiration or serum samples) and 32 were environmental samples (water or soil samples).

For clinical samples, DNA was extracted using a Maxwell instrument (Promega, Charbonnières-Les-Bains, France) according to the manufacturer’s instructions for kit CSC Blood DNA kit (reference AS1321). For water samples, DNA was extracted using BioRad Aquadien™ Bacterial DNA Extraction and Purification Kit according to the manufacturer’s instructions. Finally, for soil sample, DNA extraction was performed using the MACHEREY-NAGEL NucleoSpin Soil Kit, following manufacturer’s instruction.

To optimize and validate bioinformatic parameters, an additional set of 20 culture-positive specimens was included. DNA was extracted directly from the specimen as described above for according to samples type. In parallel, DNA was extracted from corresponding cultured isolates using the protocol described above for bacterial strains.

### Amplification of the 23S-5S by PCR

The 23S–5S ribosomal intergenic spacer region was amplified using primers previously described by Grattard et al. (2006), with minor modifications for compatibility with NGS. Each PCR reaction (final volume: 25µL) contained 10µL of Q5 ® Reaction Buffer (5X, New England Biolabs), 2.5µL of forward and reverse primer (10µM), 1 µL of dNTP (10mM), 0.5µL of Q5® High-Fidelity DNA Polymerase and 5 µL of template DNA. The amplification program included an initial denaturation step of 30 s at 98°C followed by 40 cycles of denaturation (15 s at 98 °C), annealing (20 s at 58 °C) and extension (30 s at 72 °C) and a final elongation step of 2 min at 72°C. The amplified products were migrated on an agarose gel 2% stained with SyberSafe. Samples yielding visible amplicons were subjected to sequencing.

### Library preparation and amplicon sequencing

Sequencing libraries were prepared from either genomic DNA or unpurified 23S–5S amplicons using the Illumina DNA Prep library preparation kit (Illumina®, Nextera DNA Flex Library Prep), according to the manufacturer’s protocol.

Paired-end 2□×□ 150b sequencing was performed using NextSeq 550 (Illumina®). Raw sequencing data were deposited in the European Nucleotide Archive (ENA) under study accession number ERP194788 or PRJEB114485. Individual accession numbers are listed in Supplementary Material Table S2.

### Reference database

Sequences from all assigned Legionellaceae species in NCBI taxonomy were used to manually extract the 23S-5S region. The curated reference database is publicly available at: https://github.com/genepii/nextflow_pipeline_legio/tree/main/Pipeline_qiime2_amplicons/assets.

### In-house BLASTn pipeline

Raw fastq were trimmed and assembled before sequence identification using BLASTn (Camacho et al., 2009) against the *Legionella*-specific curated database. The analysis was performed using a 95% identity threshold and 90% coverage using an in-house script available at: https://github.com/genepii/nextflow_pipeline_legio/tree/main/Pipeline_blast_amplicons. Each of the 155 reference strain-derived sequences was assessed to confirm correct species-level assignment and to exclude cross-species misclassification that could suggest multiple operon copies or ambiguous alignments.

### QIIME2 pipeline

FASTQ files were quality-trimmed using FastP version 1.3.2 (Chen, 2025) to remove Illumina adapters and discard low-quality and short reads (quality score < 30, read length < 100 bp). The resulting trimmed reads were subsequently processed with QIIME2 version 2024.10 (Bolyen et al., 2019). Paired end reads were imported into QIIME2 via *qiime tools import* using the parameter *–type ‘SampleData[PairedEndSequencesWithQuality]’*. Denoising, quality filtering, and chimera removal were performed using DADA2 (Callahan et al., 2016) (*qiime dada2 denoise*). Amplicon sequence variants (ASVs) were generated for downstream taxonomic assignment.

Taxonomic assignment was performed using two alternative QIIME2 feature-classifier approaches. First, reads were classified using VSEARCH: *qiime feature-classifier classify-consensus-vsearch*. This method searches each query sequence against the full reference database and assigns taxonomy based on a consensus of the best matching hits (up to *maxaccepts*) that share the same classification. Unlike BLAST-based approaches, it does not use local alignment scoring but relies on heuristic sequence similarity to identify matches across the entire database before selecting the best hits. We set the percentage identity to 0.95 with or without adding a percentage of coverage of 0.90.

Second, a QIIME2 Naïve Bayes classifier was trained on a curated 23S–5S database and applied for taxonomic assignment (*qiime feature-classifier fit-classifier-naive-bayes* and *classify-sklearn)*. This probabilistic approach assigns taxonomy based on the likelihood of sequence features belonging to each taxonomic group. Different confidence thresholds (0.70 to 0.99, with 0.70 as default) were tested to evaluate the trade-off between classification sensitivity and specificity.

ASV summary tables were obtained using *qiime feature-table summarize* and taxonomic composition was visualized using *qiime taxa barplot*. A threshold of 50 reads was set to consider a given species assignation.

The full bioinformatic workflow is publicly available at: https://github.com/genepii/nextflow_pipeline_legio/tree/main/Pipeline_qiime2_amplicons.

### Statistical analyses

Statistical analysis and visualization were performed in R v.3.6.1 (dplyr version 2.5.0, tidyr version 1.3.1, ggplot2 version 4.0.3).

## Results

### Specificity of the 23s-5s region

To evaluate the specificity of the 23S–5S amplicon and the curated reference database, the in-house BLASTn taxonomic assignment was performed on 155 genome-derived sequences using a 95% identity threshold and 90% coverage. All strains were correctly assigned to their respective species. Importantly, no cross-species misassignments were observed, and no evidence suggested ambiguous classification due to multiple 23S–5S operon copies within individual genomes. These results confirm the high specificity of the 23S–5S region for species-level discrimination within the genus *Legionella*.

### Optimization of Taxonomic Assignment Parameters

We then evaluated different taxonomic classification strategies using the 20 culture-positive samples to determine optimal parameters for species-level assignment.

The VSEARCH approach resulted in the assignment of three additional species in *L. pneumophila*-positive samples that were not identified by culture (*L. longbeachae, L. shakespearei*, and *L. brunensis*). Co-infection between *L. longbeachae* and *L. micdadei* was correctly identified. We further assessed the impact of applying a minimum query coverage threshold of 90% in combination with a 95% identity threshold. The addition of a coverage filter did not affect the results (Figure 1A vs 1B; Supplementary Table S3).

**Figure 1:**
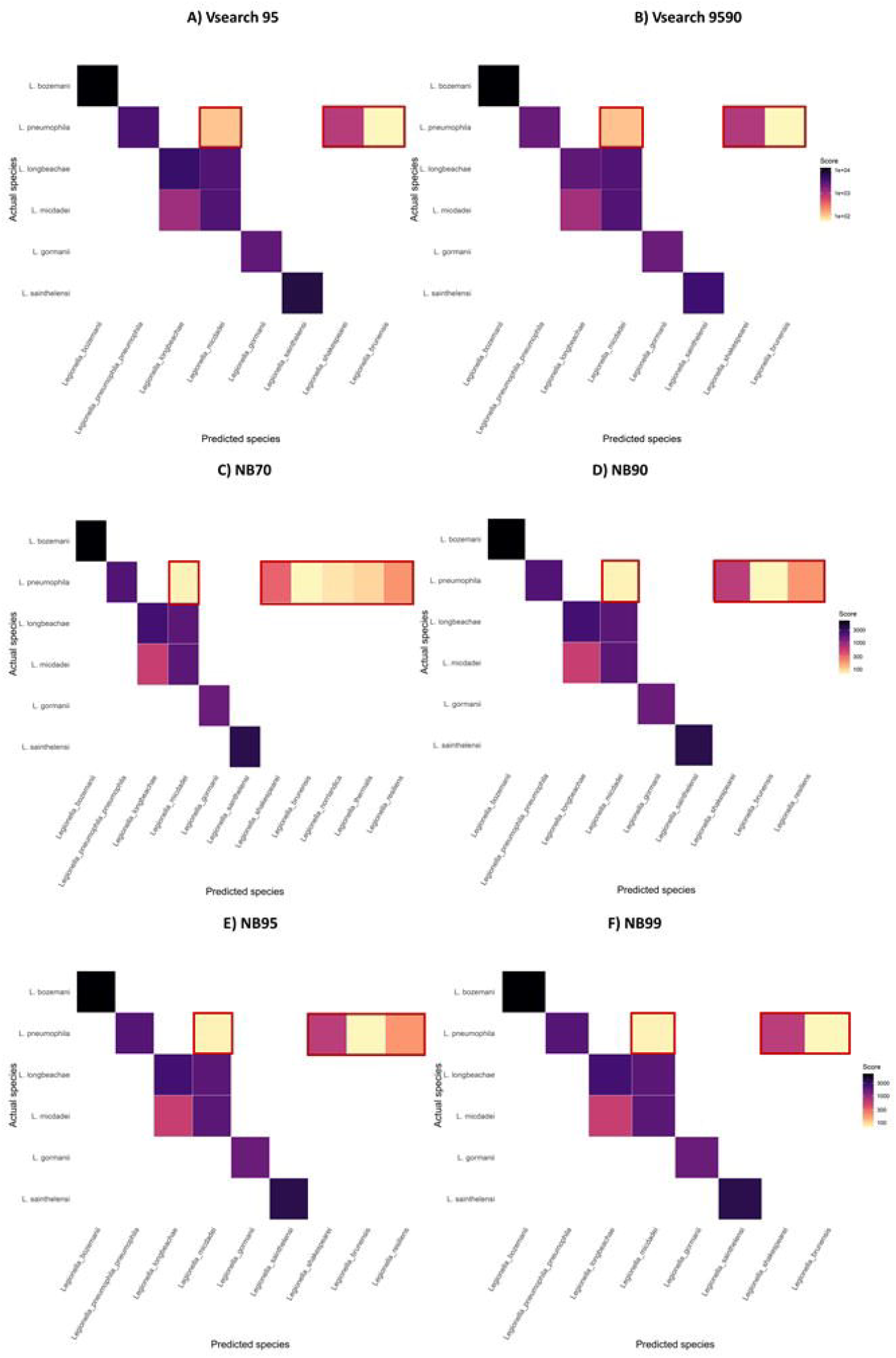
Heatmap of mean read abundance weighted by sample prevalence for species predicted by 23S-5S analyses across identified species by culture for different assignment parameters A) ≥95% VSEARCH identity, B) ≥95% VSEARCH identity and ≥90% coverage and Naïve Bayes classifier with C) 70% confidence (function default), D) 90% confidence, E) 95% confidence and F) 99% confidence. Each tile represents a species pair for which at least one sample tested positive. Tile color reflects a score defined as the mean number of reads assigned to the predicted species * (Number of samples harboring this species / Total number of samples included in the analysis). Darker colors indicate higher scores (log□ □ scale) and only species pairs with a score exceeding score of 50 are shown. Pairs absent from the plot were either not detected or fell below this threshold. Red squares show discordances between the species identified by culture and assigned species.

In contrast, the use of Naïve Bayes classifier at a confidence threshold of 70% resulted in erroneous taxonomic assignments of up to six species in *L. pneumophila*-positive samples (*L. micdadei, L. shakespearei, L. brunensis, L. norrlandica, L. thermalis and L. resiliens*,). This number decreased to four species at confidence levels of 90% and 95%, and to three species (*L. micdadei, L. shakespearei*, and *L. brunensis)* at a 99% confidence threshold (Figure 1). Co-infection between *L. longbeachae* and *L. micdadei* was correctly identified for all confidence thresholds.

As *L. shakespearei* and *L. brunensis* were consistently detected by all analytical approaches at high confidence levels, including under stringent parameter settings, we cannot exclude the possibility that these findings reflect co-infections not detected by culture-based methods.

Overall, species-level assignment was achieved in 90% (18/20) of culture-positive samples using the Naïve Bayes classifier method with a 99% confidence threshold (Table 2). As this parameter set also reduced the number of erroneous assignments, it selected for downstream analysis of culture-negative samples.

**Table 2:** Species assignation using Naïve Bayes classifier (confidence 99% level) to 23S-5S amplicons from culture-positive clinical and environmental samples from 2023 to 2025.

| Culture-positive | 2023 |  | 2024 |  |  | 2025 |  |  | TOTAL |
| --- | --- | --- | --- | --- | --- | --- | --- | --- | --- |
|  | Respiratory | Total | Respiratory | Soil | Total | Respiratory | Soil | Total |  |
| Unassigned | 0 | 0 | 0 | 1 | 1 | 1 | 0 | 1 | 2 |
| L. spp assigned | 1 | 1 | 4 | 2 | 6 | 10 | 1 | 11 | 18 |
| Total | 1 | 1 | 4 | 3 | 7 | 11 | 1 | 12 | 20 |

### Species Identification in Culture-Negative Clinical and Environmental Samples

The optimized pipeline was then applied to 92 PCR-positive, culture-negative samples collected between 2023 and 2025. Species-level assignment was achieved in 60% (55/92) of tested samples (Table 3). Among the assigned samples, *L. pneumophila* was the most frequently detected species, identified in 53% (29/55) of samples presenting with at least one read classified with a 99% confidence level (Figure 2; Supplementary Table S4).

**Table 3:** Species assignation using Naïve Bayes classifier (confidence 99% level) to 23S-5S amplicons from culture-negative clinical and environmental samples from 2023 to 2025.

|  | 2023 |  |  |  |  |
| --- | --- | --- | --- | --- | --- |
|  | Biopsy | Respiratory | Soil | Water | Total 2023 |
| Unassigned | 1 | 12 | 3 | 3 | 19 |
| L. spp assigned | 1 | 3 | 6 | 5 | 15 |
| Total | 2 | 15 | 9 | 7 | 33 |
|  | 2024 |  |  |  | Total 2024 |
|  | Biopsy | Respiratory | Serum | Soil |  |
| Unassigned | 0 | 6 | 1 | 0 | 7 |
| L. spp assigned | 1 | 9 | 0 | 3 | 13 |
| Total | 1 | 15 | 1 | 3 | 20 |
|  | 2025 |  |  |  | TOTAL 2023-2025 |
|  | Respiratory | Soil | Water | Total 2025 |  |
| Unassigned | 9 | 1 | 2 | 12 | 38 |
| L. spp assigned | 17 | 10 | 0 | 27 | 55 |
| Total | 26 | 11 | 2 | 39 | 92 |

**Figure 2.**
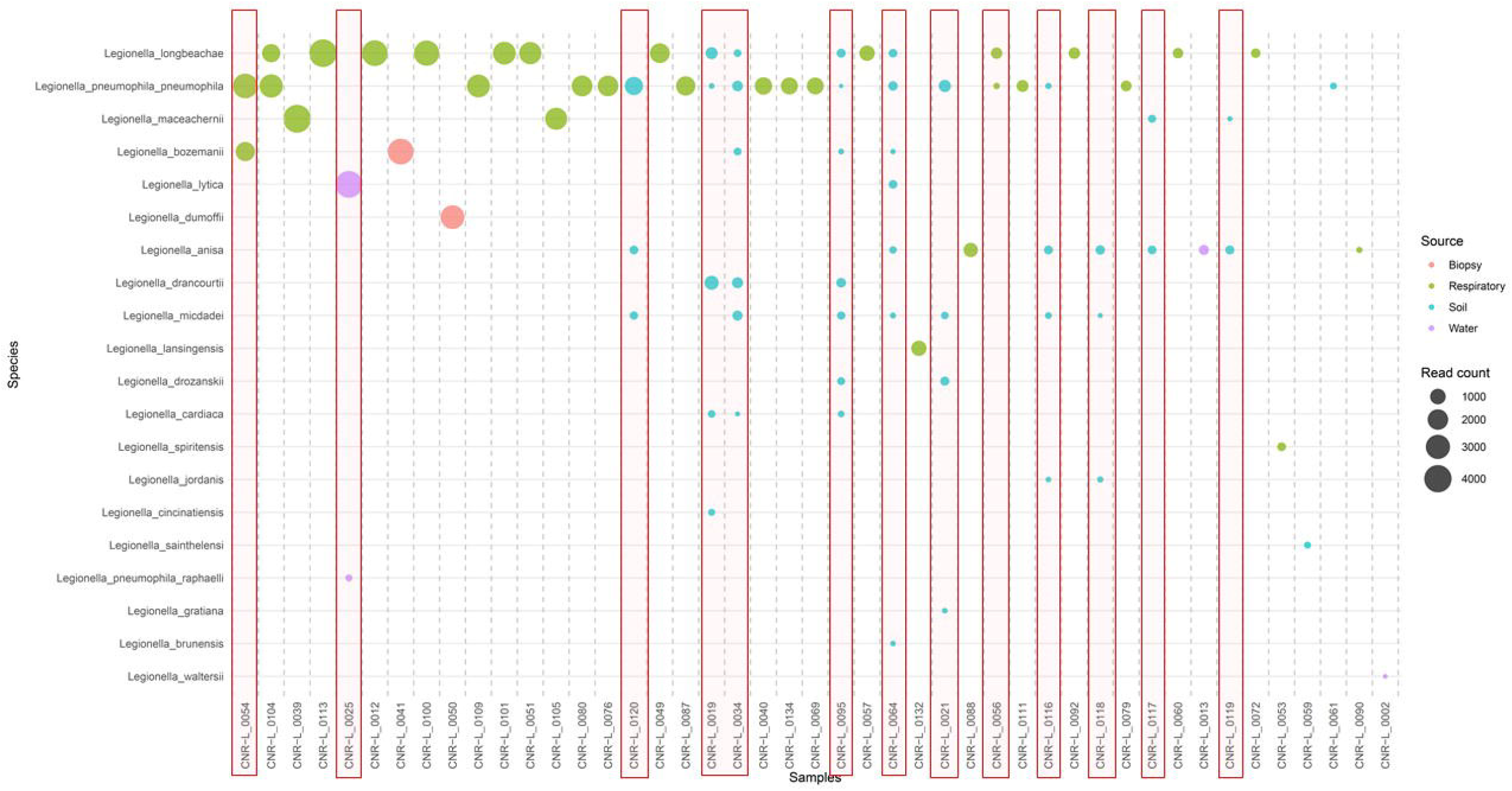
Bubble plot of *Legionella* spp. read abundance per sample detected by 23S-5S analyses using Naïve Bayes classifier (confidence 99% level). Each bubble represents a species–sample combination, with bubble size proportional to the number of reads assigned to that species (log□ □ scale) and color indicating the sample source. Species (y-axis) and samples (x-axis) are ordered by total read abundance, from lowest to highest and highest to lowest, respectively. Only species–sample combinations with at least 50 assigned reads are shown. Read squares show when multiple species of *Legionella* were detected.

As NGS technology allows the identification of multiple populations, we were able to observe that clinical and environmental specimen showed evidence of *Legionella* spp. co-occurrence. Two respiratory samples contained reads compatible with both *L. pneumophila* and *L. longbeachae*, while another respiratory sample contained reads corresponding to *L. pneumophila* and *L. bozemanii*. Environmental samples, particularly soil specimens, demonstrated substantially higher complexity. Between one and seven distinct *Legionella* spp. were detected within individual soil samples, highlighting the diversity and potential co-occurrence of multiple species in environmental reservoirs as well as the added value of NGS approaches.

## Discussion

In the present study, we developed and validated an optimized workflow for the characterization of *Legionella* spp. directly from PCR-positive, culture-negative samples using 23S-5S amplicon NGS. This approach was coupled with a curated reference database and a standardized QIIME2-based bioinformatic pipeline. Our results demonstrate that this strategy enables robust species-level identification, including in samples where several *Legionella* spp. co-occur, thereby substantially enhancing the resolution of molecular surveillance beyond conventional genus-level detection. Among the taxonomic assignment methods evaluated, the Naïve Bayes classifier achieved the highest performance, consistent with previous reports on bacterial taxonomic classification using QIIME2 and the SILVA reference database (Bokulich et al., 2018) showing that precision scores were found to be maximized when employing the Naïve Bayes classifier with stringent confidence thresholds. This confirms the suitability of the method for high-resolution *Legionella* spp. assignment in clinical and environmental samples.

Compared with alternative molecular methods, such as real-time PCR assays that are often limited to genus-level detection or to a restricted number of species targets, our approach provides broader taxonomic coverage. It also represents a complementary strategy to conventional culture, which remains limited by low sensitivity, prolonged incubation times, and dependence on bacterial viability. Furthermore, culture media are not always equally suitable for all *Legionella* spp., potentially contributing to underrepresentation of *L*. non-*pneumophila* species in routine surveillance. For example, some media, such as MWY and GVPC, support *L. micdadei* growth, whereas BMPA inhibits it due to the presence of cefamandole (Lee et al., 1993). Using the method described here, species-level assignment was achieved in 60% of PCR-positive, culture-negative samples tested, highlighting its added value in cases where isolation fails.

This NGS approach appears particularly relevant for complex environmental matrices such as soil, compost, or wastewater, where high microbial background and competing organisms frequently hinder culture-based detection even after acidification processes. The identification of up to seven *Legionella* spp. within single environmental samples underscores the ecological diversity of these reservoirs and supports the hypothesis that multiple species may co-occur in contamination sources. Importantly, we also detected co-infections in clinical specimens, including combinations of *L. pneumophila* with other species. Such findings support the added value of NGS compared to Sanger sequencing, to identify multiple populations. It may also have implications for source attribution and outbreak investigations, as routine workflows typically rely on whole-genome sequencing of a single cultured colony, potentially overlooking mixed infections or co-contamination events.

One important limitation of this approach relates to variation in the number of 23S-5S rRNA operon copies among *Legionella* spp. Although copy number in this genus ranges from one to four - narrower than in many other bacterial taxa - differences in operon count could influence relative read abundance and complicate quantitative interpretation (Hassler et al., 2022). Nevertheless, specificity testing across 155 genomes, including 90 reference genomes, demonstrated accurate species assignment without cross-species misclassification, supporting the robustness of the curated database and the selected identity threshold. However, the composition of the database, while curated to include the last described species, might also introduce bias as it may not reflect the actual diversity of *Legionella* spp. Collective effort might be needed to ensure that the database is exhaustive and kept up-to-date.

Another limitation concerns read length constraints inherent to short-read sequencing and fragmentation. The 23S-5S intergenic spacer region varies in size (approximately 210–300 bp), and in some cases, 2 × 150 bp reads may not fully overlap, potentially affecting read merging and classification efficiency especially for species with longer fragment. Longer read sequencing technologies without fragmentation such as Illumina MiSeq 2x300 or Oxford Nanopore or PacBio could address this limitation by enabling sequencing of longer ribosomal regions, thereby increasing phylogenetic resolution. Using long-read technologies also open opportunities to use even longer rDNA segments. Long-reads could also account for all operon copies. Previous studies have shown that longer ribosomal operon fragments provide enhanced discriminatory power (de Oliveira Martins et al., 2019). In anticipation of such developments, we tested the adaptability our pipeline to MiSeq and long-read data and obtained satisfactory results (data not shown).

Although robust, the proposed workflow remains relatively time-consuming and requires sequencing infrastructure and bioinformatic expertise. Alternative species-specific or multiplex PCR assays have been developed, including recent assays targeting the 23S-5S region of *L. micdadei* in environmental samples (Bélanger et al., 2025) or multiplex real-time PCR panels capable of detecting up to nine selected non-*pneumophila* species (Benitez & Winchell, 2016). However, these approaches are typically restricted to predefined species and may not perform optimally in culture-negative specimens. In contrast, the amplicon NGS strategy described here enables broad, database-driven species identification without prior assumptions regarding species composition.

The hypervariable 23S-5S ribosomal intergenic region enables species-level identification of *Legionella* directly from clinical and environmental specimens in the absence of cultured isolates. Using a curated and comprehensive reference database combined with a standardized QIIME2-based pipeline, we demonstrate high specificity and substantial diagnostic added value in PCR-positive, culture-negative samples. This study aligns with the general trend that advocate for molecular diagnostics in *Legionella* surveillance and to optimize strain-free methods for epidemiological investigation (Eble et al., 2021). Short-read 23S–5S amplicon NGS provides a practical and adaptable framework for highly-contaminated and environmental samples to improve detection of non-*pneumophila* species, identify co-infections, and enhance source attribution in outbreak settings. Integration of this method alongside conventional culture may strengthen public health surveillance, improve monitory of environmental reservoirs, and ultimately improve our understanding of the epidemiology of non-*pneumophila* Legionnaires’ disease.

## Authors statement

## Acknowledgments

The authors would like to thank the technical team of the French National Reference Center.

## Conflict of interest

The authors declare that there are no conflicts of interest.

## CRediT author contributions

**Camille Jacqueline**: Conceptualization, Methodology, Investigation, Formal analysis, Data curation, Visualization, Writing – original draft.

**Aurélie Peticca**: Investigation, Data curation, Validation, Writing – review & editing.

**Justine Lannes**: Investigation, Resources.

**Matthieu Curtil-Dit-Galin**: Writing – review & editing.

**Marine Ibranosyan**: Writing – review & editing.

**Laetitia Béraud**: Writing – review & editing.

**Ghislaine Descours**: Writing – review & editing.

**Sophie Jarraud**: Funding acquisition, Supervision, Writing – review & editing.

**Christophe Ginevra**: Conceptualization, Methodology, Supervision, Funding acquisition, Writing – review & editing.

## Funding statement

HCL are designated as EU Reference Laboratory for Public Health on *Legionella* (EURL-PH-LEGI). This publication is funded by the EU4Health programme under grant agreement 101194818, as part of the project EURL-PH-LEGI. However, views and opinions expressed are those of the authors only and do not necessarily reflect those of the European Union or the European Health and Digital Executive Agency. Neither the European Union nor the granting authority can be held responsible.

## Ethical statement

The patient specimens were submitted for *Legionella* testing, including culture from microbiology laboratories in France. This study was performed using residual anonymized clinical specimens. No additional samples were collected specifically for this study, and no identifiable patient information was accessed. According to institutional and national regulations, formal ethical approval and informed consent were not required.

## Supplementary Materials

**Table S1: ENA accession IDs for reference strains**

**Table S2: Overview of Illumina sequencing data, quality metrics, and ENA accession IDs**

**Table S3: Sample metadata and *Legionella* spp. read counts obtained by 23S-5S analyses on PCR-positive culture-positive samples**. Each sheet represents the different pipeline parameters tested. Each row corresponds to an individual sample. Columns report: the year of reception (YearReception); the sample type (Source: respiratory or water) and the culture result (Culture). The remaining columns report the number of reads assigned to each *Legionella* spp.

**Table S4: Sample metadata and *Legionella* spp. read counts obtained by 23S-5S analyses using Naïve Bayes classifier (confidence 99% level) on PCR-positive culture-negative samples**. Each row corresponds to an individual sample. Columns report: the year of reception (YearReception); the sample type (Source: respiratory or water); and the culture result (Culture). The remaining columns report the number of reads assigned to each *Legionella* spp.

## Notes

### Competing Interest Statement

The authors have declared no competing interest.

